# Cancer cells depend on environmental lipids for proliferation when electron acceptors are limited

**DOI:** 10.1101/2020.06.08.134890

**Authors:** Zhaoqi Li, Brian W. Ji, Purushottam D. Dixit, Evan C. Lien, Konstantine Tchourine, Aaron M. Hosios, Keene L. Abbott, Anna M. Westermark, Elizabeth F. Gorodetsky, Lucas B. Sullivan, Matthew G. Vander Heiden, Dennis Vitkup

## Abstract

It is not well understood how physiological environmental conditions and nutrient availability influence cancer cell proliferation. Production of oxidized biomass, which requires regeneration of the cofactor NAD+, can limit cancer cell proliferation^1-5^. However, it is currently unclear which specific metabolic processes are constrained by electron acceptor availability, and how they affect cell proliferation. Here, we use computational and experimental approaches to demonstrate that *de novo* lipid biosynthesis can impose an increased demand for NAD+ in proliferating cancer cells. While some cancer cells and tumors synthesize a substantial fraction of their lipids *de novo*^6^, we find that environmental lipids are crucial for proliferation in hypoxia or when the mitochondrial electron transport chain is inhibited. Surprisingly, we also find that even the reductive glutamine carboxylation pathway to produce fatty acids is impaired when cancer cells are limited for NAD+. Furthermore, gene expression analysis of 34 heterogeneous tumor types shows that lipid biosynthesis is strongly and consistently negatively correlated with hypoxia, whereas expression of genes involved in lipid uptake is positively correlated with hypoxia. These results demonstrate that electron acceptor availability and access to environmental lipids can play an important role in determining whether cancer cells engage in *de novo* lipogenesis to support proliferation.

## Introduction

Many proliferating cancer cells incorporate biomass precursors such as amino acids, nucleic acids, and lipids from their environment^7-10^. Nonetheless, nutrient and oxygen availability vary across different tumors^11-13^ and can promote metabolic adaptation to support cancer cell and tumor proliferation^11,14,15^. Oxidation of nutrients to produce some important biomass precursors requires regeneration of the cofactor NAD+ to serve as an electron acceptor. Recent work has demonstrated that when the mitochondrial electron transport chain (ETC) is compromised, production of oxidized biomass can be limiting for cell proliferation *in vivo* and *in vitro*^2-5,16-19^. NAD+ regeneration is important for the biosynthesis of several intermediates proliferating cells need to grow including aspartate, nucleotides, and serine^4,17,18,20,21^. However, a systems-level quantitative analysis of the oxidative power (NAD+) required for production of various biomass components and their relationship to cell proliferation is currently missing.

## Results

To better understand the potential roles of NAD+ availability and regeneration in cancer cell proliferation, we used a genome-scale stoichiometric model of human cell metabolism^22^. We applied Flux Balance Analysis (FBA)^23^ to calculate the minimal levels of NAD+ consumption required for the *de novo* synthesis of major biomass components (see full Methods). To that end, we first pruned the original genome-scale model to retain key metabolic reactions that are likely involved in biomass production^24,25^. The automatic pruning procedure was constrained such that the resulting models reproduced experimentally measured metabolic exchange fluxes for each of the NCI60 cell lines^9^. Each of the resulting FBA models comprised ∼600 reactions that reflected cell line heterogeneity and allowed the models to support biomass synthesis. We then used the obtained FBA models to estimate the minimum level of NAD+ consumption needed for the production of lipogenic acetyl-CoA from glucose and glutamine, and the synthesis of aspartate, serine, and nucleotides (see full Methods). In mammalian cells, lipogenic acetyl-CoA is reported to be synthesized through two major pathways (Fig. 1a): (1) glucose oxidation^7,26-28^, which requires NAD+ for glycolysis, the pyruvate dehydrogenase (PDH) reaction, and the TCA cycle, and (2) reductive carboxylation of glutamine-derived alpha-ketoglutarate (αKG), with production of αKG from glutamine-derived glutamate involving an oxidation reaction^1,17,26,29,30^. Notably, the computational analysis predicted that *de novo* synthesis of lipogenic acetyl-CoA, both from glucose and from glutamine, incurred a substantial NAD+ consumption cost that could exceed the demand for production of other major biomass precursors (Fig. 1a).

**Fig. 1.**
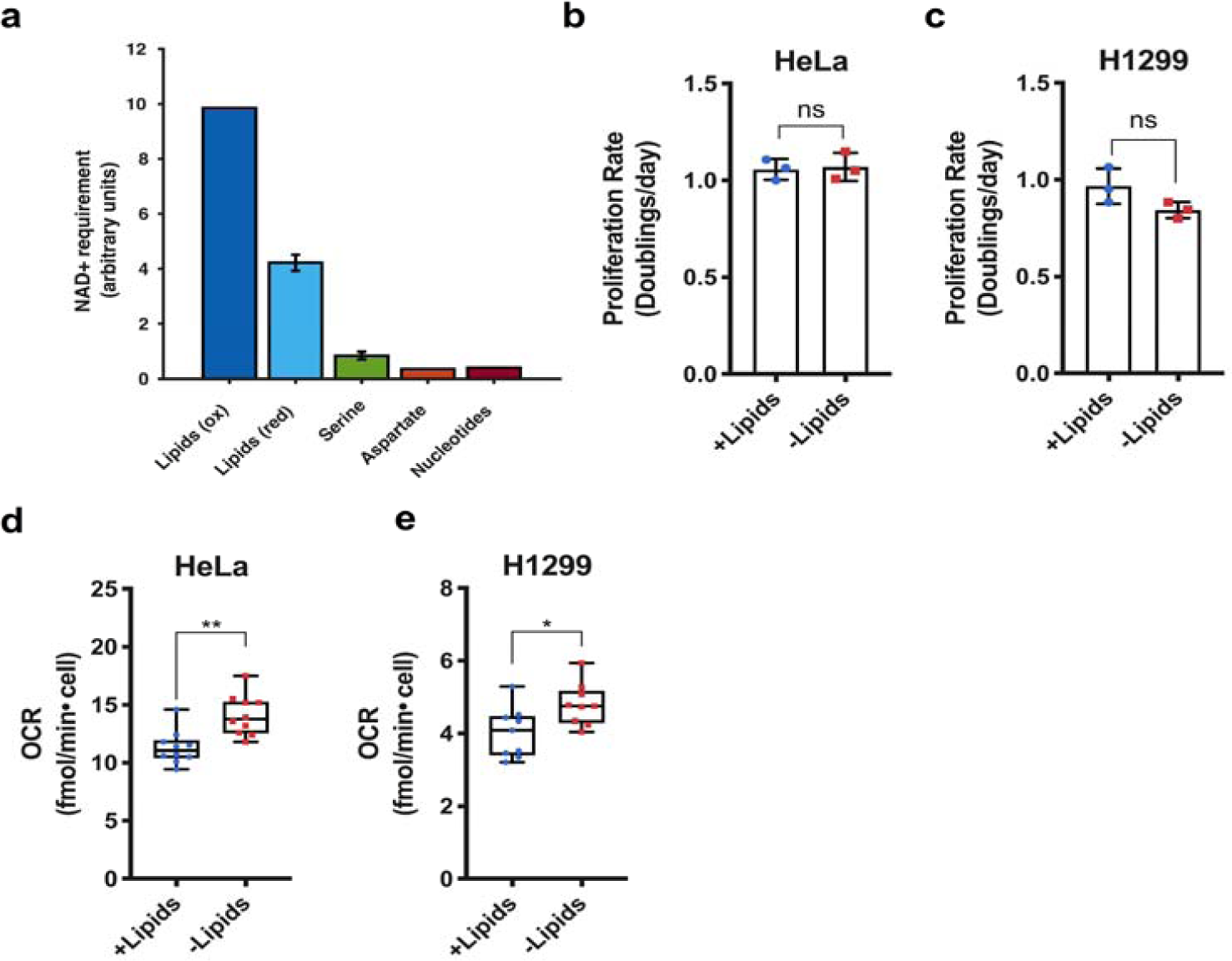
Increased lipid synthesis results in increased oxygen consumption and is predicted to increase cellular demand for NAD+. **a**, Quantitative predictions for the NAD+ consumption cost of *de novo* synthesis of specific metabolites based on metabolic flux modelling. The NAD+ consumption cost is shown for the oxidative glucose pathway (ox) and the reductive glutamine carboxylation pathway (red). Error bars denote the standard deviations of costs calculated across models of NCI-60 cancer cell lines. **b**, Cell culture media was prepared with delipidated serum, and then reconstituted with exogenous lipids (+lipids) or vehicle (–lipids). Proliferation rate of HeLa cells cultured in media +lipids or –lipids as indicated (n=3 per condition from a representative experiment). **c**, Proliferation rate of H1299 cells cultured in media +lipids or – lipids as indicated (n=3 per condition from a representative experiment). **d**, Oxygen consumption rate (OCR) of HeLa cells cultured in media +lipids or –lipids as indicated (n=3 per condition from a representative experiment). **e**, OCR of H1299 cells cultured in media +lipids or –lipids as indicated (n=3 per condition from a representative experiment). All data represent mean ± s.d. \**P* < 0.05, \*\**P* < 0.01, ns *P* > 0.05, unpaired Student’s *t*-test. All experiments were repeated 3 times or more.

To test the model predictions, we first examined how *de novo* lipid synthesis affects the rate of NAD+ regeneration. To that end, we grew cancer cells in standard normoxic tissue culture conditions with and without access to exogenous lipids^7,31^. Proliferation rates were generally similar in the presence or absence of exogenous lipids, indicating that *de novo* lipogenesis can be sufficient to support cell proliferation (Fig. 1b-c)^7^. We next confirmed that cells cultured in the absence of exogenous lipids had an increased rate of palmitate synthesis and an increased sensitivity to fatty acid synthase (FASN) inhibition (Extended Data Fig. 1). Because proliferation in lipid-deprived conditions requires upregulation of *de novo* fatty acid synthesis, we reasoned that cells would need to increase flux through reactions that regenerate NAD+. Indeed, the mitochondrial oxygen consumption rate, which reflects the rate of NAD+ regeneration by the mitochondrial ETC, increased in cells cultured in lipid-depleted conditions (Fig. 1d-e).

Our model calculations suggested that cancer cells carrying out *de novo* lipogenesis may be sensitive to NAD+ availability and thus to the reduction in the rate of NAD+ regeneration. To test this hypothesis, we considered the relationship between *de novo* lipid synthesis and hypoxia. Hypoxia has been established as a physiologically important parameter that can affect tumor growth *in vivo.* Hypoxic conditions reduce NAD+ regeneration and the NAD+/NADH ratio through decreased mitochondrial ETC activity^11,14,32^. Interestingly, culturing a diverse array of cells in the absence of exogenous lipids generally resulted in decreased proliferation rates in hypoxic but not normoxic conditions (Fig. 2a, Extended Data Fig. 2a). Supporting the idea that *de novo* lipid synthesis is impaired when electron acceptors become limiting, similar effects were also observed when cells were exposed to mitochondrial ETC inhibitors. Specifically, culturing cells with phenformin, a mitochondrial Complex I inhibitor^33^, resulted in decreased palmitate synthesis and proliferation (Fig. 2b, Extended Data Fig. 1b, Extended Data Fig. 2b,c). Culturing cells with other ETC inhibitors, such as rotenone and antimycin A, had similar effects (Fig. 2c, Extended Data Fig. 2b,c). Taken together, these data support the computational model prediction that a deficit of NAD+ can be a significant metabolic bottleneck for *de novo* lipid synthesis, and therefore can be limiting for cancer cell proliferation in the absence of exogenous lipids.

**Fig. 2.**
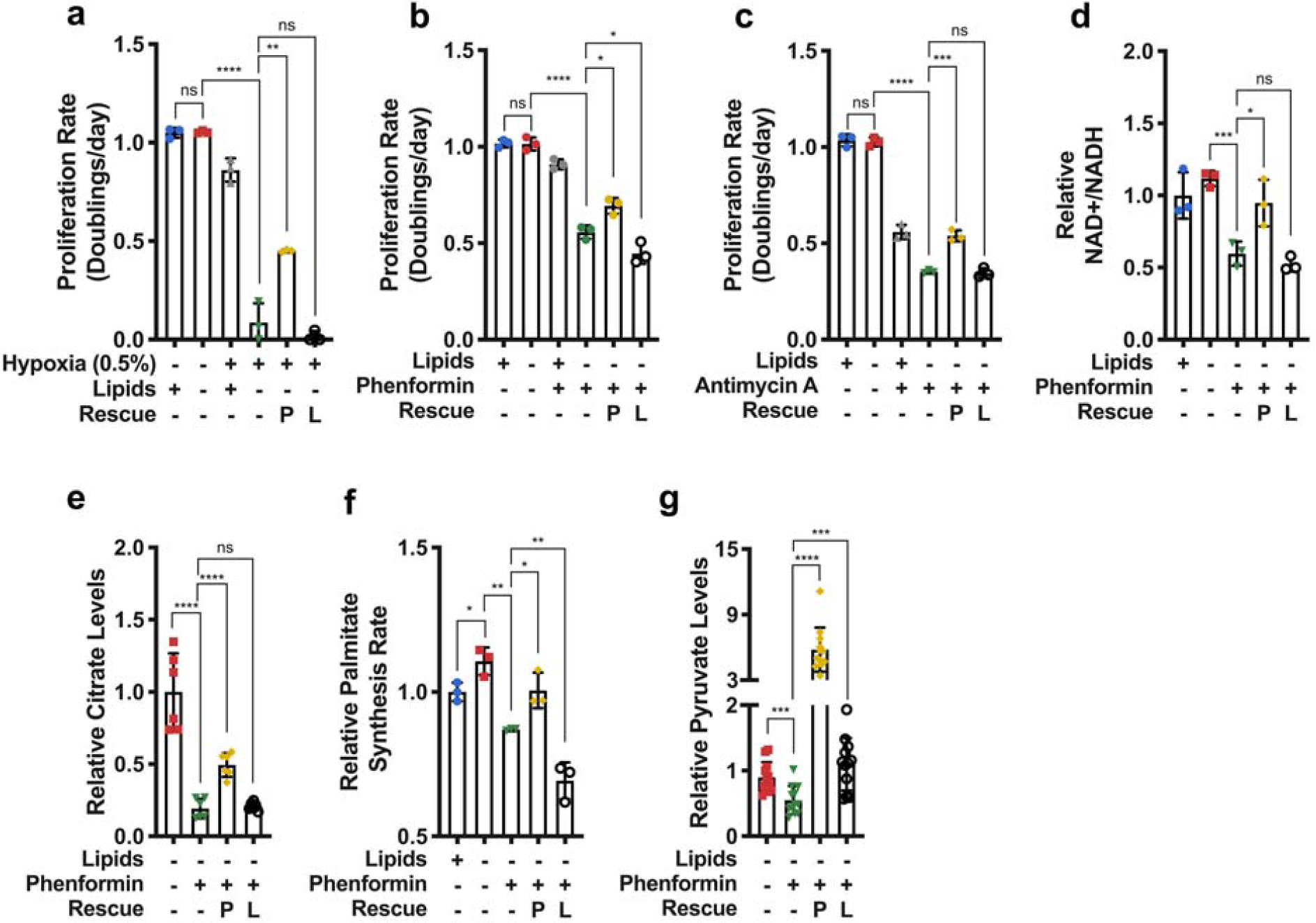
Electron acceptor availability dictates proliferation rate in the absence of exogenous lipids. **a**, Proliferation rates of HeLa cells cultured in media +lipids or –lipids in normoxia (21% oxygen) or hypoxia (0.5% oxygen), without or with pyruvate (1mM, P) and/or lactate (10mM, L) as indicated (n=3 per condition from a representative experiment). **b**, Proliferation rates of HeLa cells cultured in media +lipids or –lipids, without or with phenformin (100µM), pyruvate (1mM, P), and/or lactate (10mM, L) as indicated (n=3 per condition from a representative experiment). **c**, Proliferation rates of HeLa cells cultured in media +lipids or –lipids, without or with antimycin A (200nM), pyruvate (1mM, P), and/or lactate (10mM, L) as indicated (n=3 per condition from a representative experiment). **d**, Relative NAD+/NADH ratio in HeLa cells cultured in media +lipids or –lipids, without or with phenformin (100µM), pyruvate (1mM, P), and/or lactate (10mM, L) as indicated (n=3 per condition from a representative experiment). **e**, Relative intracellular citrate levels in HeLa cells cultured in media +lipids or –lipids, without or with phenformin (100µM), pyruvate (1mM, P), and/or lactate (10mM, L) as indicated (n=3 per condition from a representative experiment). **f**, Relative palmitate synthesis rates of HeLa cells cultured in media +lipids or –lipids, without or with phenformin (100µM), pyruvate (1mM, P), and/or lactate (10mM, L) as indicated (n=3 per condition from a representative experiment). **g**, Relative intracellular pyruvate levels in HeLa cells cultured in media +lipids or –lipids, without or with phenformin (100µM), pyruvate (1mM, P), and/or lactate (10mM, L) as indicated (n=3 per condition from a representative experiment). All data represent mean ± s.d. \**P* < 0.05, \*\**P* < 0.01, \*\*\**P* < 0.001, \*\*\*\**P* < 0.0001, ns *P* > 0.05, unpaired Student’s *t*-test. All experiments were repeated 3 times or more.

If *de novo* lipogenesis is limited by rates of NAD+ regeneration in hypoxia or under ETC inhibition, we reasoned that providing an alternative electron acceptor such as pyruvate would rescue cell proliferation in delipidated media, as exogenous pyruvate can be converted into lactate to regenerate NAD+^34,35^. Indeed, we found that pyruvate increased the NAD+/NADH ratio and the proliferation rate of cells exposed to mitochondrial ETC inhibitors (Fig. 2d, Extended Data Fig. 2f, Fig. 2b,c, Extended Data Fig. 2d-e) as well as the proliferation rate of cells in hypoxia (Fig. 2a, Extended Data Fig. 2a). Pyruvate addition also increased intracellular citrate levels (Fig. 2e) and the rate of palmitate synthesis (Fig. 2f). Furthermore, orthogonal methods of increasing the NAD+/NADH ratio, such as supplementation with α-ketobutyrate, an alternative electron acceptor^17,36^, or expression of the *L. brevis* NOX enzyme (*lb*NOX), which regenerates NAD+ by direct transfer of electrons to oxygen^5^, also rescued proliferation of phenformin-treated cells cultured in the absence of exogenous lipids (Extended Data Fig. 2g-h). These results suggest that beyond the availability of molecular oxygen^37^, cells require a functional mitochondrial ETC or alternative pathways for NAD+ regeneration to synthesize fatty acids.

Synthesis of lipids from citrate-derived acetyl-coA also requires ATP hydrolysis for the ATP-citrate lyase (ACLY) and acetyl-CoA carboxylase (ACC) reactions^38^, providing an alternative explanation for decreased lipid synthesis rates in hypoxia or under ETC inhibition, and raising the possibility that pyruvate is fueling ATP generation through the TCA cycle to rescue proliferation under these conditions^26^. To explore this possibility, we treated cells in delipidated media with phenformin in the presence or absence of exogenous lactate. While lactate can be metabolized by cells^39,40^ and provide carbons for citrate production, it also shifts the equilibrium of the lactate dehydrogenase (LDH) reaction toward pyruvate formation at the expense of consuming NAD+^16,34,35,41^. Importantly, while lactate significantly increased intracellular levels of pyruvate (Fig. 2g), it did not increase the NAD+/NADH ratio, intracellular levels of citrate, rates of palmitate synthesis, or cell proliferation (Fig. 2a-f, Extended Data Fig. 2a,d-f). These results suggest that lipid synthesis is not limited by ATP generation or carbon under these conditions. Notably, neither pyruvate nor lactate had any effect on ACC phosphorylation (Extended Data Fig. 3), a marker of increased AMP-activated protein kinase activity that is upregulated in cells experiencing energy stress^42^.

To better understand the specific metabolic pathways that support *de novo* fatty acid synthesis and how they may be shaped by electron acceptor availability, we next performed kinetic isotope tracing experiments to assess the metabolite fates of U-^13^C-Glucose or U-^13^C-Glutamine. We found that culturing cells in the absence of exogenous lipids resulted in increased citrate synthesis from both glucose and glutamine via oxidative pathways as measured by the rate of 2-carbon-labeled citrate (M+2) formation from U-^13^C-Glucose, or the rate of 4-carbon-labeled citrate (M+4) formation from U-^13^C-Glutamine (Fig. 3a-c). These results suggest that cancer cells may utilize pathways with high NAD+ requirements for fatty acid synthesis when electron acceptors are available. As expected, the increased oxidative metabolism was accompanied by an increase in fatty acid synthase (FASN) and mitochondrial pyruvate carrier 2 (MPC2) expression (Fig. 3d). We also observed a decrease in the inhibitory phosphorylation of serine 293 on the E1/A subunit of the PDH complex^43,44^, which catalyzes pyruvate conversion to acetyl-CoA and mediates the entry of pyruvate into the TCA cycle^45-49^ (Fig. 3d).

**Fig. 3.**
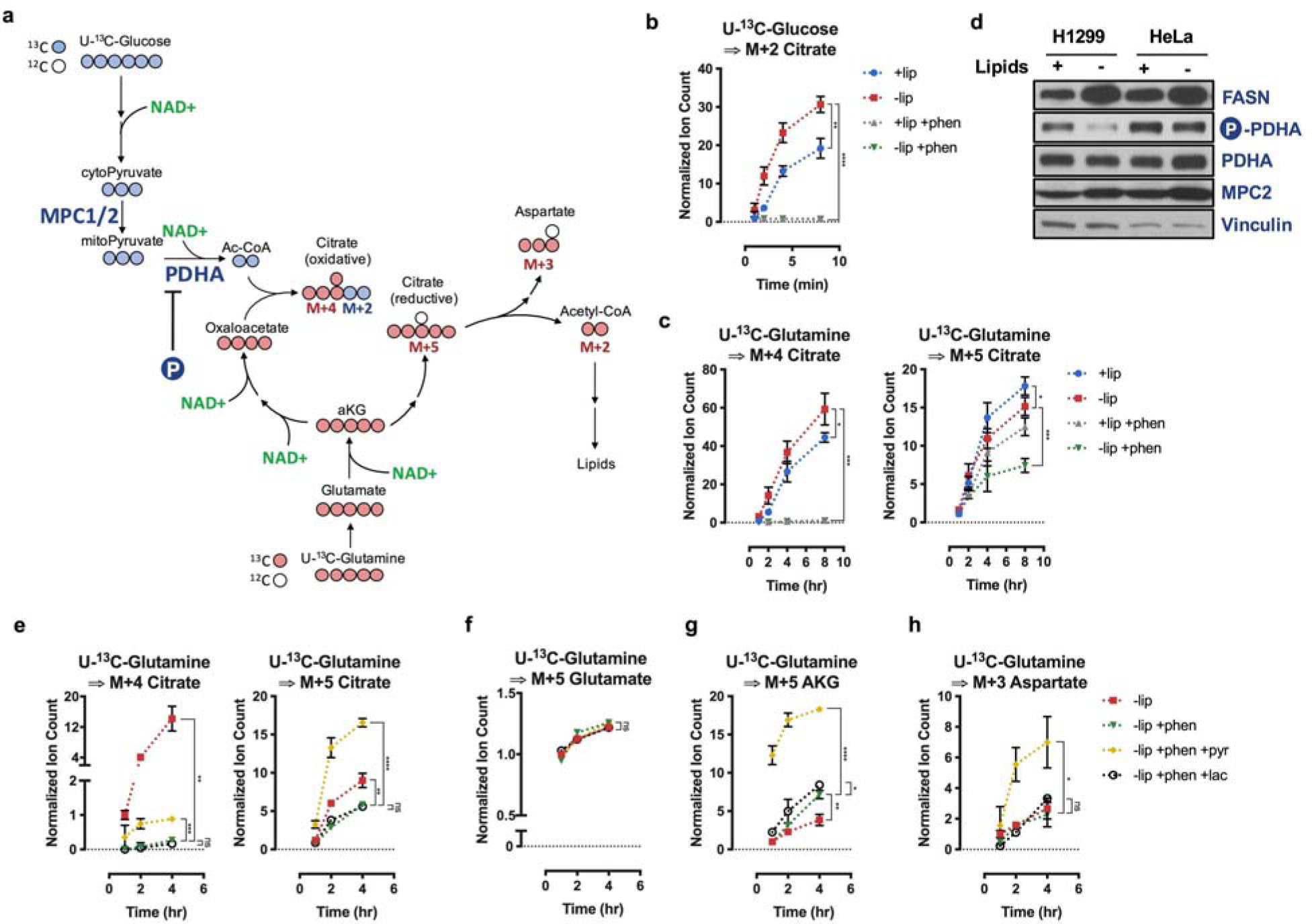
Electron Reductive TCA cycle flux is gated by electron acceptor availability. **a**, Schematic depicting carbon transitions involved in the oxidative and reductive pathways to generate citrate from glucose and glutamine. Oxidation of glucose carbons requires mitochondrial pyruvate entry, mediated through MPC1/2, as well as PDHA activity, which is negatively regulated through phosphorylation. Oxidatively produced citrate will be labeled on two carbons (M+2) from U-^13^C-Glucose or four carbons (M+4) from U-^13^C-Glutamine. Reductively produced citrate will be labeled on five carbons (M+5) from U-^13^C-Glutamine; cleavage of M+5 citrate will yield three carbon labeled oxaloacetate (M+3), which is in equilibrium with M+3 aspartate. **b**, Assessment of oxidative metabolism via production of M+2 citrate from U-^13^C-Glucose over time in HeLa cells cultured in media +lipids or –lipids, with and without phenformin (100µM) as indicated. Data were normalized to M+2 citrate in +lipids at t=2min (n=3 per condition from a representative experiment). **c**, Assessment of oxidative and reductive glutamine metabolism to produce M+4 and M+5 citrate from U-^13^C-Glutamine over time in HeLa cells cultured media +lipids or –lipids, with and without phenformin (100µM) as indicated. Data were normalized to M+4 citrate in +lipids at t=1hr (n=3 per condition from a representative experiment). **d**, Immunoblot analysis of FASN expression, PHDA serine 293 phosphorylation, total PDHA expression, and MPC2 expression in HeLa and H1299 cells cultured in media +lipids or –lipids as indicated. Vinculin was used as a loading control. **e**, Assessment of M+4 and M+5 citrate production from U-^13^C-Glutamine over time in HeLa cells cultured in media –lipids, without or with phenformin (100µM), without or with pyruvate (1mM, pyr) or lactate (10mM, lac) as indicated. Data were normalized to M+4 citrate in –lipids at t=1hr (n=3 per condition from a representative experiment). **f**, Assessment of M+5 glutamate production over time from U-^13^C-Glutamine in HeLa cells cultured in media –lipids without or with phenformin (100µM), without or with pyruvate (1mM, pyr) or lactate (10mM, lac) as indicated. Data were normalized to M+5 glutamate in –lipids at t=1hr (n=3 per condition from a representative experiment). **g**, Assessment of M+5 αKG production over time from U-^13^C-Glutamine in HeLa cells cultured in media –lipids without or with phenformin (100µM), without or with pyruvate (1mM, pyr) or lactate (10mM, lac) as indicated. Data were normalized to M+5 αKG in –lipids at t=1hr (n=3 per condition from a representative experiment). **h**, Assessment of M+3 aspartate production over time from U-^13^C-Glutamine in HeLa cells cultured in media – lipids without or with phenformin (100µM), without or with pyruvate (1mM, pyr) or lactate (10mM, lac) as indicated. Data were normalized to (M+3) aspartate in –lipids at t=1hr (n=3 per condition from a representative experiment). All data represent mean ± s.d. \**P* < 0.05, \*\**P* < 0.01, \*\*\**P* < 0.001, \*\*\*\**P* < 0.0001, ns *P* > 0.05, unpaired Student’s *t*-test. All experiments were repeated 3 times or more.

Prior work has shown that hypoxia and pharmacological ETC inhibition result in a decreased fractional contribution of glucose carbon to lipogenic citrate, and a concomitantly increased fractional contribution of glutamine carbon via reductive carboxylation of glutamine-derived αKG^50-52^. In agreement with these findings, culturing cells with phenformin in the presence or absence of exogenous lipids resulted in decreased M+4 citrate labeling and increased M+5 citrate labeling from U-^13^C-Glutamine (Extended Data Fig. 4a). These changes were also accompanied by an increase in the αKG/citrate ratio (Extended Data Fig. 4b), also consistent with previous observations^53,54^. Moreover, the increased relative contribution of glutamine carbon to produce M+5 citrate was also accompanied by a decrease in total citrate levels (Extended Data Fig. 4c). Surprisingly, using kinetic tracing we found that the rates of both oxidative and reductive glutamine flux substantially decreased in cells treated with phenformin (Fig. 3c) compared to untreated cells, suggesting that reductive carboxylation does not fully compensate for a decrease in citrate synthesis from glucose when NAD+ is limited. These data support the results of the computational analysis that both oxidative and reductive pathways of lipogenic citrate synthesis incur a substantial NAD+ consumption cost, and that both routes may be compromised when the NAD+/NADH ratio decreases (Fig. 1a).

Given that pyruvate was able to increase the rate of fatty acid synthesis by regenerating NAD+, we next sought to determine the mechanism through which this was achieved. We therefore traced U-^13^C-Glutamine and assessed the rate of M+4 and M+5 citrate formation (Fig. 3e). We observed that in cells cultured in delipidated media with phenformin, the addition of pyruvate, but not lactate, increased rates of citrate production through both oxidative and reductive routes (Fig. 3e). However, the rate of glutamine reductive carboxylation far exceeded the rate of glucose/glutamine oxidation (Fig. 3e), likely because the NAD+ requirement for reductive carboxylation is lower than for the oxidative route (Fig. 1a). Importantly, the increased rate of reductive carboxylation of glutamine-derived carbon appeared to be facilitated by an increase in the rate of αKG production from glutamate (Fig. 3g) and not from increased glutamate production from glutamine (Fig. 3f). As the production of αKG from glutamate converts a carbon-nitrogen bond on glutamate into a carbon-oxygen double bond on αKG, and can occur via either glutamate dehydrogenase or transamination reactions^1,17^, these results are consistent with the conversion of glutamate to αKG being net oxidative and therefore dependent on the NAD+/NADH ratio. We also confirmed that the citrate produced via reductive glutamine flux was further metabolized by ACLY as the rate of M+3 aspartate appearance from U-^13^C-glutamine was elevated in the pyruvate-treated, but not lactate-treated cells (Fig. 3h). Overall, these experiments suggest that the carboxylation of glutamine-derived αKG to produce fatty acids, a net reductive pathway, may be substantially inhibited due to reactions affected by a decreased NAD+/NADH ratio, and that this bottleneck can impair cell proliferation.

Exogenous acetate can be metabolized by cancer cells in culture and by tumors^55,56^. Previous studies have shown that acetate can have various context-dependent metabolic fates, ranging from its usage in epigenetic modifications to serving as a source of carbon for lipid synthesis^55,57-59^. We rationalized that acetate would circumvent the NAD+ cost of lipid synthesis by directly contributing to cytosolic acetyl-CoA, and thus support cell proliferation in electron acceptor limited conditions^60^. Indeed, exogenous acetate rescued proliferation of cells cultured in lipid-depleted conditions in the presence of phenformin or antimycin A (Fig. 4a-b, Extended Data Fig. 5a-b), without altering the intracellular NAD+/NADH ratio (Fig. 4c, Extended Data Fig 5c). Exogenous acetate also restored the proliferation of lipid-deprived cells in hypoxia (Fig. 4g, Extended Data Fig. 5d). Consistent with its role as a direct carbon source for fatty acid synthesis, exogenous acetate increased the palmitate synthesis rate (Fig. 4d) without changing citrate levels (Fig. 4e), TCA cycle fluxes (Fig. 4f), levels of TCA cycle intermediates (Extended Data Fig. 6a-d) or ACC phosphorylation (Extended Data Fig. 3). These data are consistent with acetate restoring fatty acid synthesis by circumventing the NAD+ requirement for citrate production. Notably, the conversion of exogenous acetate to lipogenic acetyl-CoA consumes ATP^61^, further supporting a model of lipid biosynthesis being limited by the NAD+/NADH ratio rather than ATP availability under these conditions.

**Fig. 4.**
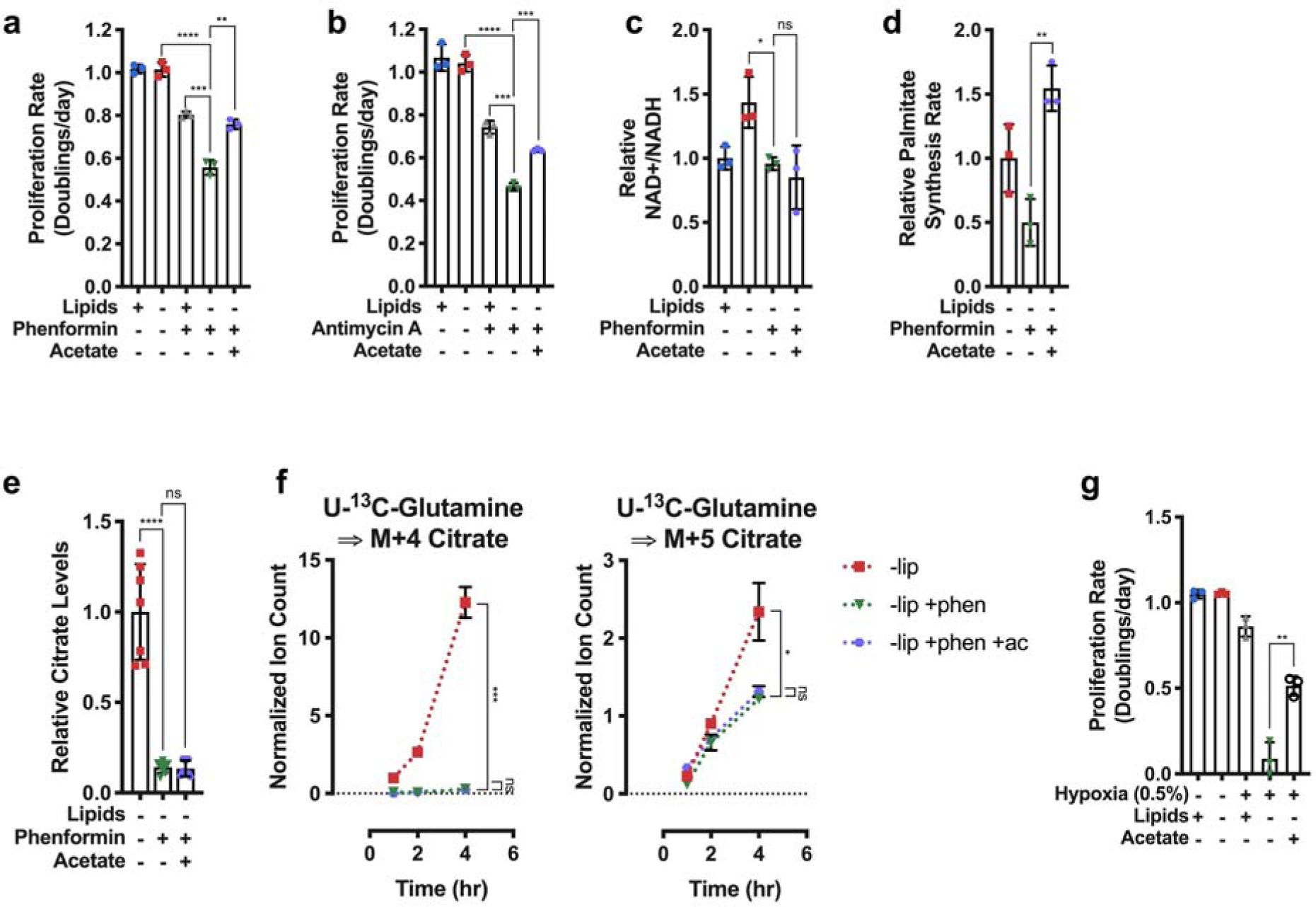
Bypassing oxidative steps in fatty acid synthesis rescues proliferation in electron acceptor deficient cells. **a**, Proliferation rates of HeLa cells cultured in media +lipids or –lipids, without or with phenformin (100µM) or acetate (200 µM) as indicated (n=3 per condition from a representative experiment). **b**, Proliferation rates of HeLa cells cultured in media +lipids or – lipids, without or with antimycin A (200nM) or acetate (200 µM) as indicated (n=3 per condition from a representative experiment). **c**, Relative NAD+/NADH ratio in HeLa cells cultured in media +lipids or –lipids, without or with phenformin (100µM) or acetate (200 µM) as indicated (n=3 per condition from a representative experiment). **d**, Relative palmitate synthesis rates of HeLa cells cultured in media –lipids without or with phenformin (100µM) or acetate (200 µM) as indicated (n=3 per condition from a representative experiment). **e**, Relative intracellular citrate levels in HeLa cells cultured in media –lipids without or with phenformin (100µM) or acetate (200 µM) as indicated (n=3 per condition from a representative experiment). **f**, Assessment of M+4 and M+5 citrate production from U-^13^C-Glutamine over time in HeLa cells cultured in media –lipids without or with phenformin (100µM) or acetate (200 µM) as indicated. Data were normalized to M+4 citrate in –lipids at t=1hr (n=3 per condition from a representative experiment). **g**, Proliferation rates of HeLa cells cultured in media +lipids or –lipids, in normoxia (21% oxygen) or hypoxia (0.5% oxygen), and without or with acetate (200 µM) as indicated. Data from the first four conditions are the same as those presented in Fig. 2a. (n=3 per condition). All data represent mean ± s.d. \**P* < 0.05, \*\**P* < 0.01, \*\*\**P* < 0.001, \*\*\*\**P* < 0.0001, unpaired Student’s *t*-test. All experiments were repeated 3 times or more.

Our findings raise the possibility that *de novo* lipid synthesis in tumors may be affected by conditions where electron acceptors are limiting *in vivo*. We therefore asked whether hypoxia was associated with coordinated changes in metabolic gene expression across human tumors. We used RNA-sequencing data from The Cancer Genome Atlas (TCGA) to investigate the correlations between mRNA expression of 87 KEGG-defined metabolic pathways^62^ and expression signatures of known tumor hypoxia markers^32,63^. Strikingly, analysis of over 10,000 different primary tumor samples demonstrated that lipid biosynthesis was one of the most negatively correlated metabolic pathways to the tumor hypoxia expression signature (Pearson’s R = −0.39, p < 10^−16^) (Fig. 5a,b). The correlation was also significant when controlling for the potential effects of tumor proliferation rate, using purine, pyrimidine, and tRNA synthesis (partial Pearson’s correlation R = −0.39, p < 10^−16^, R = −0.38, p < 10^−16^, and R = −0.32, p < 10^−16^, respectively). Out of the 34 tumor types we investigated, 29 types individually displayed a significant negative correlation (at 1% False Discovery Rate (FDR)) between hypoxia and fatty acid biosynthesis (Extended Data Fig. 7a,b). Across all tumor samples, we also observed that the hypoxia expression signature positively correlated with the expression of genes involved in fatty acid uptake (Pearson’s R = 0.41, p < 10^−16^) (Fig. 5c); this finding was also significant in 28 out of the 34 tumor types analyzed individually (at 1% FDR, Extended Data Fig 7a,c). These results suggest that multiple tumors adapt to hypoxia by downregulating *de novo* fatty acid synthesis and instead relying more on lipid acquisition from the environment to support biomass production.

**Fig. 5.**
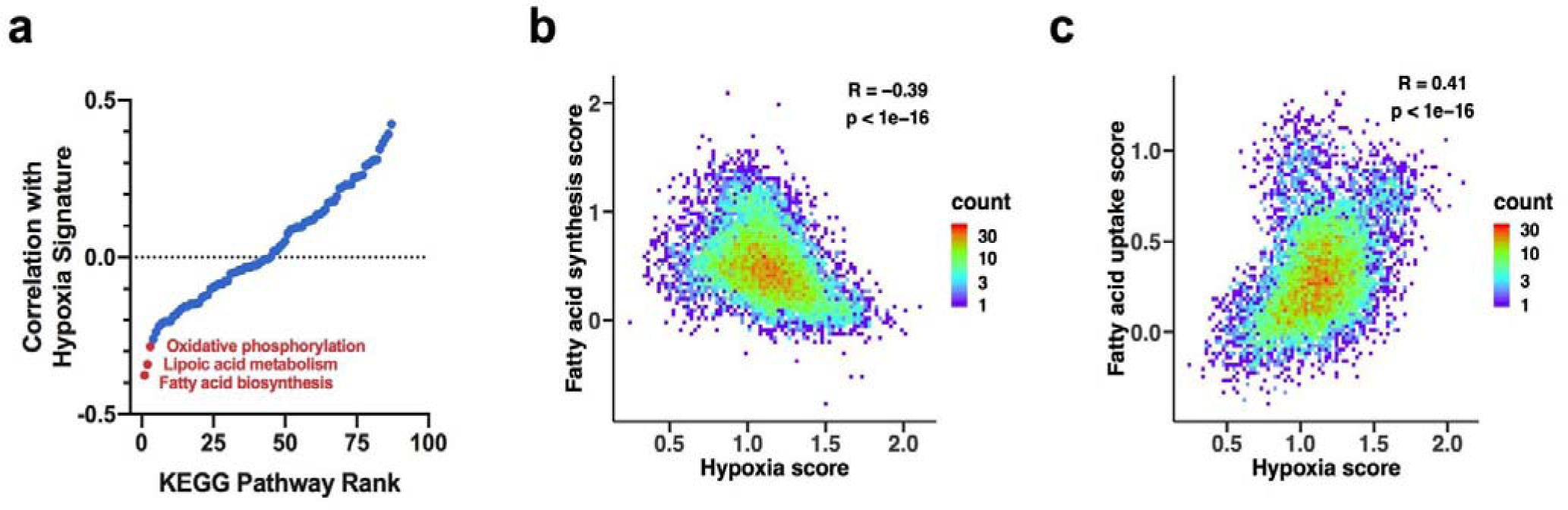
Correlations between mRNA expression of lipid synthesis and lipid uptake genes and expression of gene markers of tumor hypoxia. **a**, Ranking of 87 KEGG metabolic pathways based on the correlation between expression of their genes with expression of known markers of tumor hypoxia (tumor hypoxia scores); the correlations were calculated using TCGA expression data, such that stronger negative correlations correspond to stronger anti-correlations with hypoxia markers. **b**, Density plot of the correlation between the tumor hypoxia score and expression of genes forming the fatty acid synthesis pathway. **c**, Density plot of the correlation between the tumor hypoxia score and expression of genes participating in fatty acid uptake. In **b** and **c**, density counts represent the number of TCGA samples with the corresponding tumor hypoxia scores and fatty acid biosynthesis or fatty acid uptake gene expression levels.

## Discussion

Cancer cells optimize metabolism to support proliferation in diverse tissue and environmental contexts^11,12,15,64-66^. Environmental metabolic constrains on cancer cell proliferation include the availability of nutrients, such as exogenous amino acids and lipids, and other variables such as hypoxia. An important metabolic role of environmental oxygen is to serve as a terminal electron acceptor for the mitochondrial ETC, facilitating the regeneration of NAD+ to support oxidation reactions. Some tumors, in contrast to many normal tissues, synthesize a substantial fraction of their lipids *de novo*^6^ and cancer cells rely on reductive glutamine carboxylation for lipid synthesis in hypoxia^50-52^. Our computational and experimental analyses demonstrate that *de novo* lipids biosynthesis incurs substantial NAD+ consumption costs which can be high relative to the production costs of other biomass components. This can make *de novo* lipid synthesis, specifically lipogenic citrate production, limiting for proliferation in hypoxia or other conditions where NAD+ regeneration is impaired. This suggests exogenous lipid availability will be important for cell proliferation in NAD+-limited condition, explaining why a hypoxia-associated gene expression signature correlates with decreased expression of *de novo* lipid synthesis genes in most tumor types.

When considering both cofactors NAD+ and NADP+, the use of glutamine carbon to make palmitate, a pathway previously reported as an alternative route for lipid synthesis in hypoxia or following mitochondrial inhibition^50-52^, is chemically net reductive. Surprisingly, we find that this net reductive pathway is also impaired in the absence of exogenous electron acceptors because it is gated by the oxidative conversion of glutamate to αKG. The gating of net reductive pathways by reactions sensitive to the availability of oxidizing co-factors may also apply to pathways beyond those investigated in the present study. More broadly, our findings illustrate how electron acceptor deficits can reshape essential metabolic programs and create dependencies of cancer cell growth on specific environmental nutrient sources, such as lipids and acetate.

## Acknowledgements

We thank the members of the Vitkup and Vander Heiden lab for helpful discussions. The results published here are in part based upon data generated by the TCGA Research Network. This work is supported by the NIH grant R01CA201276 (D.V., M.G.V.H.), T32GM007367 (B.W.J.), the M.D.-Ph.D. program at Columbia University (B.W.J.), the NIH grant U54CA209997 (P.D., K.T., B.W.J., D.V.), and T32GM007287 (Z.L., K.L.A.). E.C.L. is a Damon Runyon Fellow supported by the Damon Runyon Cancer Research Foundation (DRG-2299-17). A.M.H. was supported by an HHMI International Student Fellowship. L.B.S. acknowledges support from a Pathway to Independence award from the NIH (K99CA218679/R00CA218679). E.F.G. was supported by the MIT MSRP program. M.G.V.H. is also supported by the Lustgarten Foundation, SU2C, the Ludwig Center at MIT, the NCI (R35CA242379), the MIT Center for Precision Cancer Medicine, the Emerald Foundation, and a Faculty Scholar award from HHMI.

## Contributions

B.W.J, Z.L., P.D.D., and D.V. conceived the study. Z.L., B.W.J., P.D.D., M.G.V.H. and D.V. wrote the manuscript. B.W.J., P.D.D., and K.T. developed and executed the computation analysis of global metabolic flux and NAD+ costs analysis. Z.L., A.M.H., E.F.G., K.L.A., and L.B.S. performed proliferation assays. Z.L. performed oxygen consumption assays. Z.L. and E.C.L. performed serum delipidation. Z.L. performed kinetic isotope tracing and lipid synthesis assays. Z.L. performed immunoblot assays. A.M.W. performed NAD+ measurement assays. Z.L. performed mass spectrometry and analysis for metabolites. B.W.J., P.D.D., and K.T. performed TCGA analysis of gene expression correlations. M.G.V.H. and D.V. supervised the project.

## Materials and Methods

### Cell Culture Experiments

Cell lines were cultured in RPMI (Fisher Scientific, MT10040CV) supplemented with 10% fetal bovine serum. For all experiments, cells were washed three times in phosphate buffered saline (PBS), and then cultured in 4mL of RPMI with 10% dialyzed fetal bovine serum, and supplemented with lactate, pyruvate, acetate as indicated. The reagents used for the cell culture experiments are as follows: sodium pyruvate (Sigma, P2256), sodium L-lactate (Sigma, L7022), sodium acetate (Sigma, S2889), Lipid Mixture 1, Chemically Defined (Sigma, L0288), rotenone (Sigma, R8875), phenformin hydrochloride (Sigma, P7045 FLUKA), antimycin A from *streptomyces sp.* (A8676), GSK2194069 (Tocris, 5303), U-^13^C-Glucose (Cambridge Isotope Laboratories, CLM-1396), U-^13^C-Glutamine (Cambridge Isotope Laboratories, CLM-1822), U-^13^C-Pyruvate (Cambridge Isotope Laboratories, CLM-2440), U-^13^C-Lactate (Cambridge Isotope Laboratories, CLM-1579), 99% enriched deuterium oxide (Sigma, 151882). All cells were cultured at 37°C with 5% CO_2_. When indicated, +lipid culture conditions refer to RPMI supplemented with 10% delipidated serum and 1% Lipid Mixture 1.

### Oxygen Consumption

An Agilent Seahorse Bioscience Extracellular Flux Analyzer (XF24) was used to measure oxygen consumption rates (OCR). Cells were plated at 50,000 cells per well in Seahorse Bioscience 24-well plates in 50 μL of RPMI supplemented with 10% dialyzed fetal bovine serum. An additional 500 μL of media was added following a 1 hour incubation. The following day, cells were washed three times with PBS and incubated in RPMI with the indicated treatment. Five hours later, OCR measurements were made every 6 minutes, and injections of pyruvate at 16 minutes, FCCP (Sigma, C2920) at 32 minutes, and rotenone and antimycin A at 48 minutes. After OCR data acquisition, each well of the plate was collected for cell number analysis. Basal OCR was calculated by subtracting residual OCR following the addition of rotenone and antimycin A from the initial OCR measurements.

### Proliferation Rates

Cells were plated in replicate six-well plates in 2mL at an initial seeding density of 20,000 cells. Cells were permitted to settle overnight and a six-well dish was counted to calculate the starting cell number at the initiation of the experiment. For all remaining plates, cells were washed three times with PBS and 4 mL of treatment media was added to each well. Three days after the initial treatment, cells were quantified using a sulforhdamine B (SRB) colorimetric assay. All SRB measurements are normalized to a blank. Proliferation rates were calculated using the equation described below:

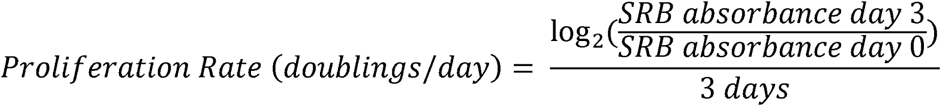

### GCMS Metabolite Measurements

Gas-chromatography coupled to mass spectrometry (GCMS) analysis was done as described previously^1^. For polar metabolites, dried samples were derivatized with 20μL of methoxamine (MOX) reagent (ThermoFisher TS-45950) and 25μL of N-tert-butyldimethylsilyl-*N*-methyltrifluoroacetamide with 1% tert-butyldimethylchlorosilane (Sigma 375934). Following derivatization, samples were analyzed using a DB-35MS column ((30m × 0.25mm i.d. × 0.25 μm, Agilent J&W Scientific) in an Agilent 7890 gas chromatograph (GC) coupled to an Agilent 5975C mass spectrometer (MS). For fatty acids, dried samples were first resuspended in 100μL toluene (Sigma, 650579) and 200uL of 2% sulfuric acid in methanol was added. Samples were incubated overnight at 50°C. The next day, 500uL 5% NaCl and 500uL hexane (Sigma, 34859) were added to the sample. The top fraction was collected, followed by a second hexane extraction with 500uL. The hexane fractions were pooled and dried down under nitrogen gas. The dried samples were resuspended in 50uL hexane and transferred into glass inserts for GCMS analysis. FAME samples were analyzed using a DB-FastFAME GC Column, 30 m x 0.25 mm x 0.25 µm, 7 in. configuration, Agilent J&W Scientific) in an Agilent 7890 gas chromatograph coupled to an Agilent 5975C mass spectrometer. The chromatograph method for fatty acid analysis was as follows: hold at 100 °C for 5 minutes, followed by a ramp of 8°C/min to 180 °C, then followed by a ramp of 1 °C/min to 230 °C. Data were analyzed and corrected for natural isotope abundance using in-house algorithms.

### Dynamic isotope tracing experiments

Cells were plated in six-well plates in at a seeding density of 150,000 cells per well. Cells were permitted to settle overnight. Prior to the initiation of the experiment, cells were washed three times with PBS, and then cultured in 2mL media containing the relevant tracers (10mM Glucose, 2mM Glutamine, 1mM Pyruvate, 10mM Lactate, or 45% enriched deuterium oxide) and the indicated treatment condition. For polar metabolite analysis, following the appropriate incubation, wells were washed as quickly as possible with ice-cold blood bank saline and lysed on the dish with 400μL of ice-cold 80% HPLC grade methanol (Sigma, 646377) in HPLC grade water (Sigma, 270733) with 1μg/400μL norvaline (Sigma, N7627) to use as an internal extraction standard. Samples were scraped, collected into Eppendorf tubes, and vortexed for 10 minutes at 4°C. Samples were centrifuged at 21,000 × g for 10 minutes at 4°C to precipitate the protein. The supernatant was dried down under nitrogen gas for subsequent analysis by GCMS. For analysis of fatty acid methyl esters (FAME), following the appropriate incubation, wells were washed three times as quickly as possible with ice-cold blood bank saline and lysed on the dish with 700μL of (4:3) methanol:0.88% KCl in water with 0.25μg/mL tridecanoic acid (Sigma, T0502) to use as an internal extraction standard. Samples were scraped, collected into glass vials (Thermo Fisher Scientific, C4010-1), and 800uL HPLC-grade dichloromethane (Thermo Fisher Scientific, 402152) was added. Samples were vortexed for 10 minutes at 4°C and centrifuged at 5,000 × g for 10 minutes at 4°C. The bottom fraction was collected into glass vials and dried down under nitrogen gas for subsequent analysis by GCMS.

### Palmitate Synthesis Rate

Cells were plated in six-well plates in at a seeding density of 150,000 cells per well. Cells were permitted to settle overnight. Prior to the initiation of the experiment, cells were washed three times with PBS, and then cultured for 24 hours in the indicated treatment condition. After 24 hours, media with the indicated treatment condition and reconstituted with 99% deuterium oxide was added to each well to achieve a final deuterium oxide enrichment of 45%. After 2 hours of culture in tracer medium, cells were harvested, and fatty acids were analyzed by FAME GCMS as described above. Palmitate synthesis rates were calculated by integrating the sum of all labeled species and normalized to cell number and time of label treatment.

### NAD+/NADH Measurements

Cells were seeded at 20,000 cells per well in six-well plates and permitted to adhere overnight. Next, cells were washed three times in PBS and incubated in 4 mL of the indicated treatment media for 5 hours. Cells were then rapidly washed three times in 4°C PBS and extracted in 100μL of ice-cold lysis buffer (1% dodecyltrimethylammonium bromide [DTAB] in 0.2 N of NaOH diluted 1:1 with PBS), snap-frozen in liquid nitrogen and frozen at −80°C. The NAD+/NADH ratio was measured using a protocol adapted from the manual of the NAD/NADH-Glo Assay kit (Promega G9072)^2^. To measure NAD+, 20μL of lysate was transferred to PCR tubes and diluted with 20μL of lysis buffer and 20μL 0.4 N HCl, and subsequently incubated at 60°C for 15 minutes. For NADH measurement, 20μL of freshly thawed lysate was transferred to PCR tubes and incubated at 75°C for 30 minutes. The acidic conditions permit for selective degradation of NADH, while the basic conditions degrade NAD+. Following the incubation, samples were spun on a bench-top centrifuge and quenched with 20μL neutralizing solution. The neutralizing solution consisted of 0.5 M Tris base for NAD+ samples and 0.25 M Tris in 0.2 N HCl for the NADH samples. The instructions in the Promega G9072 technical manual were then followed to measure NAD+ and NADH levels using a luminometer (Tecan Infinite M200Pro).

### Immunoblotting

Cells washed with ice-cold PBS, and scraped into cold RIPA buffer containing cOmplete Mini protease inhibitor (Roche 11836170001) and PhosStop Phosphatase Inhibitor Cocktail Tablets (Roche 04906845001). Protein concentration was calculated using the BCA Protein Assay (Pierce 23225) with BSA as a standard. Lysates were resolved by SDS-PAGE and proteins were transferred onto nitrocellulose membranes using the iBlot2 Dry Blotting System (Thermo Fisher, IB21001, IB23001). Protein was detected with the primary antibodies anti-Pyruvate Dehydrogenase E1-alpha subunit (phospho S293) (Abcam, ab92696), anti-Pyruvate Dehydrogenase E1-alpha subunit (total) (Proteintech, 18068-1-AP), MPC2 (Cell Signaling Technologies, 46141S), FASN (Cell Signaling Technologies, 3180S), and anti-Vinculin (Sigma, V9131). The secondary antibody used was anti-rabbit IgG HRP-linked antibody (Cell Signaling Technologies, 7074S).

## References

1. Hosios, A. M. & Vander Heiden, M. G. The redox requirements of proliferating mammalian cells. Journal of Biological Chemistry 293, 7490–7498 (2018).

2. Garcia-Bermudez, J. et al. Aspartate is a limiting metabolite for cancer cell proliferation under hypoxia and in tumours. Nature Cell Biology 20, 1–12 (2018).

3. Sullivan, L. B. et al. Aspartate is an endogenous metabolic limitation for tumour growth. Nature Cell Biology 20, 1–12 (2018).

4. Diehl, F. F., Lewis, C. A., Fiske, B. P. & Vander Heiden, M. G. Cellular redox state constrains serine synthesis and nucleotide production to impact cell proliferation. Nature Metabolism (2019).

5. Titov, D. V. et al. Complementation of mitochondrial electron transport chain by manipulation of the NAD+/NADH ratio. Science 352, 231–235 (2016).

6. Santos, C. R. & Schulze, A. Lipid metabolism in cancer. FEBS Journal 279, 2610–2623 (2012).

7. Hosios, A. M. et al. Amino acids rather than glucose account for the majority of cell mass in proliferating mammalian cells. Dev. Cell 36, 540–549 (2016).

8. Yao, C.-H. et al. Exogenous Fatty Acids Are the Preferred Source of Membrane Lipids in Proliferating Fibroblasts. Cell Chemical Biology 23, 483–493 (2016).

9. Jain, M. et al. Metabolite Profiling Identifies a Key Role for Glycine in Rapid Cancer Cell Proliferation. Science 336, 1040 (2012).

10. Damaraju, V. L. et al. Nucleoside anticancer drugs: the role of nucleoside transporters in resistance to cancer chemotherapy. Oncogene 22, 7524–7536 (2003).

11. Muir, A., Danai, L. V. & Vander Heiden, M. G. Microenvironmental regulation of cancer cell metabolism: implications for experimental design and translational studies. Disease Models & Mechanisms 11, dmm035758–12 (2018).

12. Sullivan, M. R. et al. Quantification of microenvironmental metabolites in murine cancers reveals determinants of tumor nutrient availability. Elife 8, e44235 (2019).

13. Vaupel, P., Kallinowski, F. & Okunieff, P. Blood flow, oxygen and nutrient supply, and metabolic microenvironment of human tumors: a review. Cancer Res 49, 6449 (1989).

14. Eales, K. L., Hollinshead, K. E. R. & Tennant, D. A. Hypoxia and metabolic adaptation of cancer cells. Oncogenesis 5, e190–8 (2016).

15. Hu, J. et al. Heterogeneity of tumor-induced gene expression changes in the human metabolic network. Nature Biotechnology 31, 522–529 (2013).

16. Gui, D. Y. et al. Environment Dictates Dependence on Mitochondrial Complex I for NAD+ and Aspartate Production and Determines Cancer Cell Sensitivity to Metformin. Cell Metabolism 24, 716–727 (2016).

17. Sullivan, L. B. et al. Supporting Aspartate Biosynthesis Is an Essential Function of Respiration in Proliferating Cells. Cell 162, 552–563 (2015).

18. Birsoy, K. et al. An Essential Role of the Mitochondrial Electron Transport Chain in Cell Proliferation Is to Enable Aspartate Synthesis. Cell 162, 540–551 (2015).

19. Fernandez-de-Cossio-Diaz, J. & Vazquez, A. Limits of aerobic metabolism in cancer cells. Sci. Rep. 7, 13488 (2017).

20. Bao, X. R. et al. Mitochondrial dysfunction remodels one-carbon metabolism in human cells. Elife 5, e10575 (2016).

21. Murphy, J. P. et al. The NAD+ Salvage Pathway Supports PHGDH-Driven Serine Biosynthesis. Cell Reports 24, 2381–2391.e5 (2018).

22. Duarte, N. C. et al. Global reconstruction of the human metabolic network based on genomic and bibliomic data. Proc Natl Acad Sci USA 104, 1777 (2007).

23. Orth, J. D., Thiele, I. & Palsson, B. Ø. What is flux balance analysis? Nature Biotechnology 28, 245–248 (2010).

24. Jerby, L., Shlomi, T. & Ruppin, E. Computational reconstruction of tissue-specific metabolic models: application to human liver metabolism. Mol Syst Biol 6, 401 (2010).

25. Zielinski, D. C. et al. Systems biology analysis of drivers underlying hallmarks of cancer cell metabolism. Sci. Rep. 7, 41241 (2017).

26. Saggerson, E. D. The regulation of glyceride synthesis in isolated white-fat cells. The effects of acetate, pyruvate, lactate, palmitate, electron-acceptors, uncoupling agents and oligomycin. Biochem J 128, 1069–1078 (1972).

27. Wise, E. M., Jr & Ball, E. G. Malic enzyme and Lipogenesis. Proc Natl Acad Sci USA 52, 1255–1263 (1964).

28. Liu, L. et al. Malic enzyme tracers reveal hypoxia-induced switch in adipocyte NADPH pathway usage. Nat. Chem. Biol. 12, 345–352 (2016).

29. DeBerardinis, R. J. et al. Beyond aerobic glycolysis: Transformed cells can engage in glutamine metabolism that exceeds the requirement for protein and nucleotide synthesis. Proc Natl Acad Sci USA 104, 19345 (2007).

30. Santos, C. R. & Schulze, A. Lipid metabolism in cancer. FEBS Journal 279, 2610–2623 (2012).

31. Hosios, A., Li, Z., Lien, E. & Vander Heiden, M. Preparation of Lipid-Stripped Serum for the Study of Lipid Metabolism in Cell Culture. BIO-PROTOCOL 8, 1–8 (2018).

32. Harris, A. L. Hypoxia — a key regulatory factor in tumour growth. Nature Reviews Cancer 2, 38–47 (2002).

33. Owen, M. R., Doran, E. & Halestrap, A. P. Evidence that metformin exerts its anti-diabetic effects through inhibition of complex 1 of the mitochondrial respiratory chain. Biochem J 348 Pt 3, 607–614 (2000).

34. Bücher, T. et al. State of Oxidation-Reduction and State of Binding in the Cytosolic NADH-System as Disclosed by Equilibration with Extracellular Lactate/Pyruvate in Hemoglobin-Free Perfused Rat Liver. Eur J Biochem 27, 301–317 (1972).

35. Hung, Y. P., Albeck, J. G., Tantama, M. & Yellen, G. Imaging Cytosolic NADH-NAD+ Redox State with a Genetically Encoded Fluorescent Biosensor. Cell Metabolism 14, 545–554 (2011).

36. Kim, W. et al. Polyunsaturated Fatty Acid Desaturation Is a Mechanism for Glycolytic NAD+ Recycling. Cell Metabolism 1–42 (2019).

37. Kamphorst, J. J. et al. Hypoxic and Ras-transformed cells support growth by scavenging unsaturated fatty acids from lysophospholipids. Proc Natl Acad Sci USA 110, 8882 (2013).

38. Hatzivassiliou, G. et al. ATP citrate lyase inhibition can suppress tumor cell growth. Cancer Cell 8, 311–321 (2005).

39. Faubert, B. et al. Lactate Metabolism in Human Lung Tumors. Cell 171, 358–371.e9 (2017).

40. Hui, S. et al. Glucose feeds the TCA cycle via circulating lactate. Nature 551, 115–118 (2017).

41. Zhao, Y. et al. In vivo monitoring of cellular energy metabolism using SoNar, a highly responsive sensor for NAD+/NADH redox state. Nat Protoc 11, 1345–1359 (2016).

42. Garcia, D. & Shaw, R. J. AMPK: Mechanisms of Cellular Energy Sensing and Restoration of Metabolic Balance. Mol. Cell 66, 789–800 (2017).

43. Kaplon, J. et al. A key role for mitochondrial gatekeeper pyruvate dehydrogenase in oncogene-induced senescence. Nature 498, 109–112 (2013).

44. Linn, T. C., Pettit, F. H. & Reed, L. J. α-keto acid dehydrogenase complexes, X. Regulation of the activity of pyruvate dehydrogenase complex from beef kidney mitochondria by phosphorylation and dephosphorylation. Proc Natl Acad Sci USA 62, 234 (1969).

45. Schell, J. C. et al. A Role for the Mitochondrial Pyruvate Carrier as a Repressor of the Warburg Effect and Colon Cancer Cell Growth. Mol. Cell 56, 400–413 (2014).

46. Vacanti, N. M. et al. Regulation of Substrate Utilization by the Mitochondrial Pyruvate Carrier. Mol. Cell 56, 425–435 (2014).

47. Yang, C. et al. Glutamine Oxidation Maintains the TCA Cycle and Cell Survival during Impaired Mitochondrial Pyruvate Transport. Mol. Cell 56, 414–424 (2014).

48. Bricker, D. K. et al. A Mitochondrial Pyruvate Carrier Required for Pyruvate Uptake in Yeast, Drosophila, and Humans. Science 337, 96 (2012).

49. Herzig, S. et al. Identification and Functional Expression of the Mitochondrial Pyruvate Carrier. Science 337, 93 (2012).

50. Wise, D. R. et al. Hypoxia promotes isocitrate dehydrogenase-dependent carboxylation of α-ketoglutarate to citrate to support cell growth and viability. Proc Natl Acad Sci USA 108, 19611 (2011).

51. Metallo, C. M. et al. Reductive glutamine metabolism by IDH1 mediates lipogenesis under hypoxia. Nature 481, 380–384 (2012).

52. Mullen, A. R. et al. Reductive carboxylation supports growth in tumour cells with defective mitochondria. Nature 481, 385–388 (2011).

53. Fendt, S.-M. et al. Reductive glutamine metabolism is a function of the α-ketoglutarate to citrate ratio in cells. Nature Communications 4, 2236 (2013).

54. Gameiro, P. A. et al. In Vivo HIF-Mediated Reductive Carboxylation Is Regulated by Citrate Levels and Sensitizes VHL-Deficient Cells to Glutamine Deprivation. Cell Metabolism 17, 372–385 (2013).

55. Comerford, S. A. et al. Acetate Dependence of Tumors. Cell 159, 1591–1602 (2014).

56. Mashimo, T. et al. Acetate Is a Bioenergetic Substrate for Human Glioblastoma and Brain Metastases. Cell 159, 1603–1614 (2014).

57. Schug, Z. T. et al. Acetyl-CoA synthetase 2 promotes acetate utilization and maintains cancer cell growth under metabolic stress. Cancer Cell 27, 57–71 (2015).

58. Bulusu, V. et al. Acetate Recapturing by Nuclear Acetyl-CoA Synthetase 2 Prevents Loss of Histone Acetylation during Oxygen and Serum Limitation. Cell Reports 18, 647–658 (2017).

59. Liu, X. et al. Acetate Production from Glucose and Coupling to Mitochondrial Metabolism in Mammals. Cell 175, 502–513.e13 (2018).

60. Röhrig, F. & Schulze, A. The multifaceted roles of fatty acid synthesis in cancer. Nature Reviews Cancer 16, 732–749 (2016).

61. Jones, M. E., Lipmann, F., Hilz, H. & Lynen, F. On the enzymatic mechanism of Coenzyme A acetylation with adenosine triphosphate and acetate. J. Am. Chem. Soc. 75, 3285–3286 (1953).

62. Kanehisa, M. & Goto, S. KEGG: Kyoto Encyclopedia of Genes and Genomes. Nucleic Acids Res. 28, 27–30 (2000).

63. Mense, S. M. et al. Gene expression profiling reveals the profound upregulation of hypoxia-responsive genes in primary human astrocytes. Physiological Genomics 25, 435–449 (2006).

64. Davidson, S. M. et al. Environment Impacts the Metabolic Dependencies of Ras-Driven Non-Small Cell Lung Cancer. Cell Metabolism 23, 517–528 (2016).

65. Davidson, S. M. et al. Direct evidence for cancer-cell-autonomous extracellular protein catabolism in pancreatic tumors. Nat Med 23, 235–241 (2017).

66. Sullivan, M. R. et al. Increased Serine Synthesis Provides an Advantage for Tumors Arising in Tissues Where Serine Levels Are Limiting. Cell Metabolism 29, 1410–1421.e4 (2019).

## Methods References

1. Lewis, C. A. et al. Tracing Compartmentalized NADPH Metabolism in the Cytosol and Mitochondria of Mammalian Cells. Mol. Cell 55, 253–263 (2014)

2. Gui, D. Y. et al. Environment Dictates Dependence on Mitochondrial Complex I for NAD+ and Aspartate Production and Determines Cancer Cell Sensitivity to Metformin. Cell Metabolism 24, 716–727 (2016).

