## Supplemental Figures for "Cancer cells depend on environmental lipids for proliferation when electron acceptors are limited"

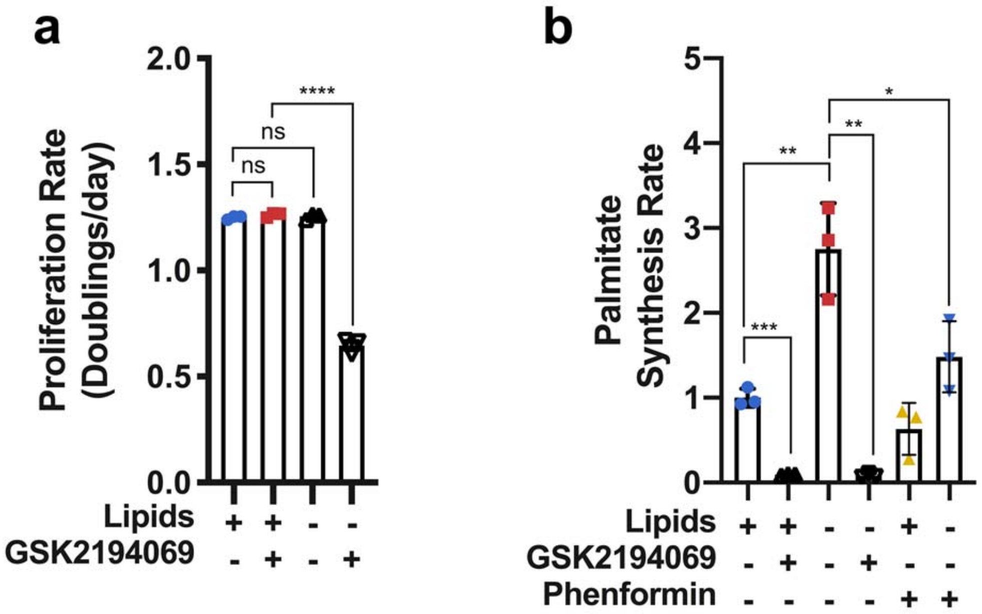


**Extended Data Fig. 1 | Effects of lipid depletion and fatty acid synthesis inhibition on cell proliferation. a,** Cell culture media was prepared with delipidated serum, and then reconstituted with exogenous lipids (+lipids) or vehicle (–lipids). Proliferation rates of HeLa cells cultured in media +lipids or –lipids without and with the FASN inhibitor GSK2194069 (0.3 μM) as indicated (n=3 per condition from a representative experiment). **b**, Relative palmitate synthesis rates of HeLa cells cultured in media +lipids or –lipids without and with GSK2194069 (0.3 μM), or with and without phenformin (100 μM) as indicated (n=3 per condition from a representative experiment). All data represent mean ± s.d. *$P$ < 0.05, **$P$ < 0.01, ***$P$ < 0.001, ****$P$ < 0.0001, ns $P$ > 0.05, unpaired Student's $t$-test. All experiments were repeated 3 times or more.

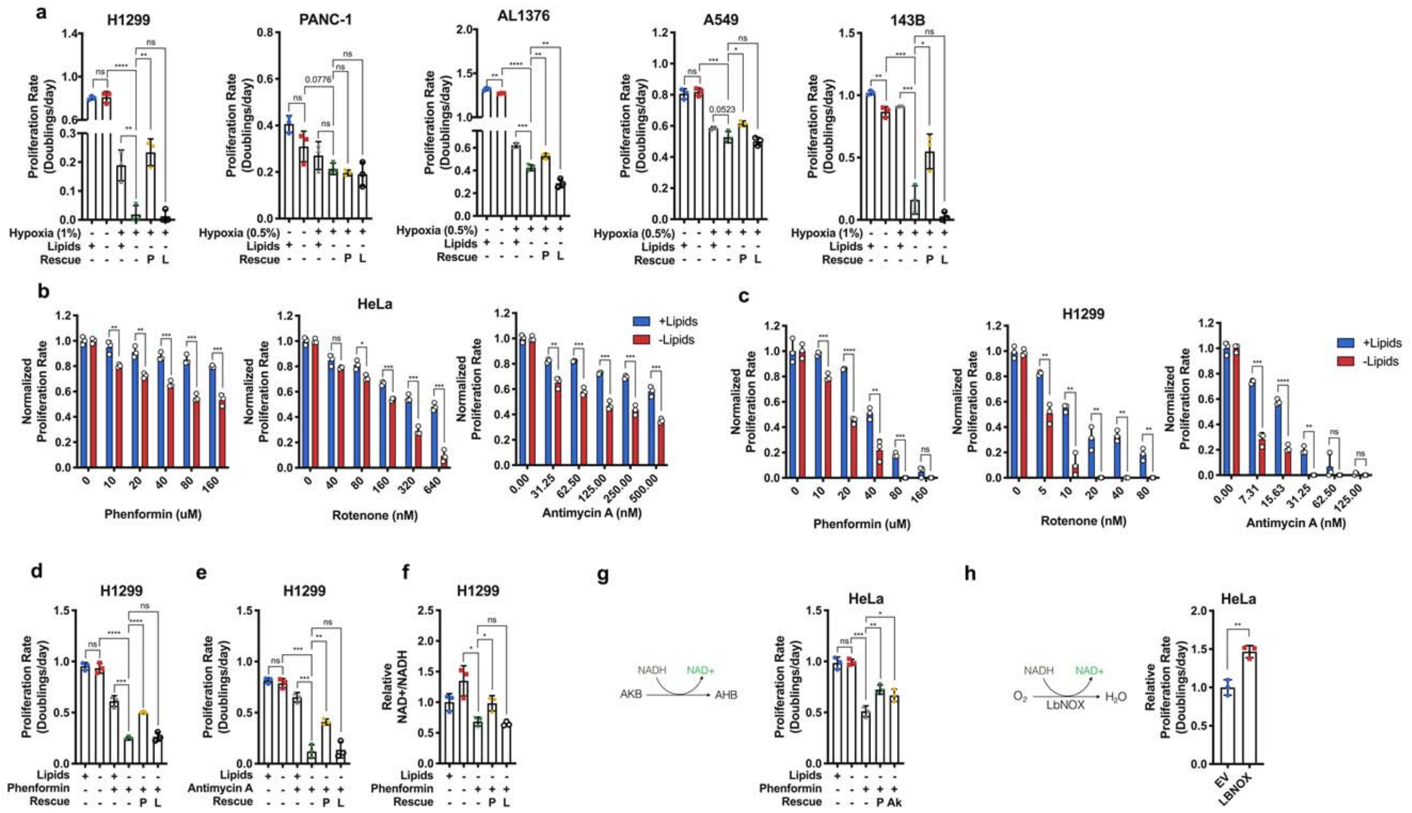


**Extended Data Fig. 2 | Electron acceptor availability dictates proliferation rate in the absence of exogenous lipids.** **a**, Proliferation rates of H1299, PANC-1, AL1376, A549, and 143B cells cultured in media +lipids or –lipids in normoxia (21% oxygen) or hypoxia (0.5% or 1% oxygen), without or with pyruvate (1mM, P) and/or lactate (10mM, L) as indicated (n=3 per condition from a representative experiment). **b**, Proliferation rate of HeLa cells cultured in media +lipids or –lipids with a titration of phenformin (Complex I inhibitor), rotenone (Complex I inhibitor), or antimycin A (Complex III inhibitor) as indicated (n=3 per condition from a representative experiment). **c**, Proliferation rate of H1299 cells cultured in media +lipids or –lipids with a titration of phenformin, rotenone, or antimycin A as indicated (n=3 per condition from a representative experiment). **d**, Proliferation rates of H1299 cells cultured in media +lipids or –lipids, without or with phenformin (10μM), pyruvate (1mM, P), and/or lactate (10mM, L) as indicated (n=3 per condition from a representative experiment). **e**, Proliferation rates of H1299 cells cultured in media +lipids or –lipids without or with antimycin A (15nM), pyruvate (1mM, P), and/or lactate (10mM, L) as indicated (n=3 per condition from a representative experiment). **f**, Relative NAD+/NADH ratio in H1299 cells cultured in media +lipids or –lipids without or with phenformin (10µM), pyruvate (1mM, P), and/or lactate (10mM, L) as indicated (n=3 per condition from a representative experiment). **g**, Proliferation rates of HeLa cells cultured in media +lipids or –lipids without or with phenformin (100µM), pyruvate (1mM, P), and/or alpha-ketobutyrate (1mM, Ak) as indicated (n=3 per condition from a representative experiment). **h**, Relative proliferation rates of HeLa cells expressing empty vector (EV) or *lb*NOX cultured in –lipids with phenformin (100µM) as indicated. Data were normalized to HeLa-EV cells (n=3 per condition from a representative experiment). All data represent mean ± s.d. *$P < 0.05$, **$P < 0.01$, ***$P < 0.001$, ns $P > 0.05$, unpaired Student's *t*-test. All experiments were repeated 3 times or more.

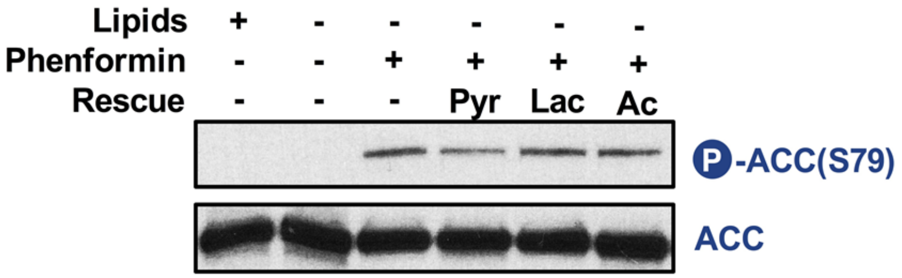


**Extended Data Fig. 3 | Effects of exogenous metabolites on ACC phosphorylation**

Immunoblot of total ACC and ACC serine 79 phosphorylation in HeLa cells cultured for 24 hours in media +lipids or –lipids without or with phenformin (100µM), pyruvate (1mM, Pyr), lactate (10mM, Lac), and/or acetate (200µM, Ac) as indicated.

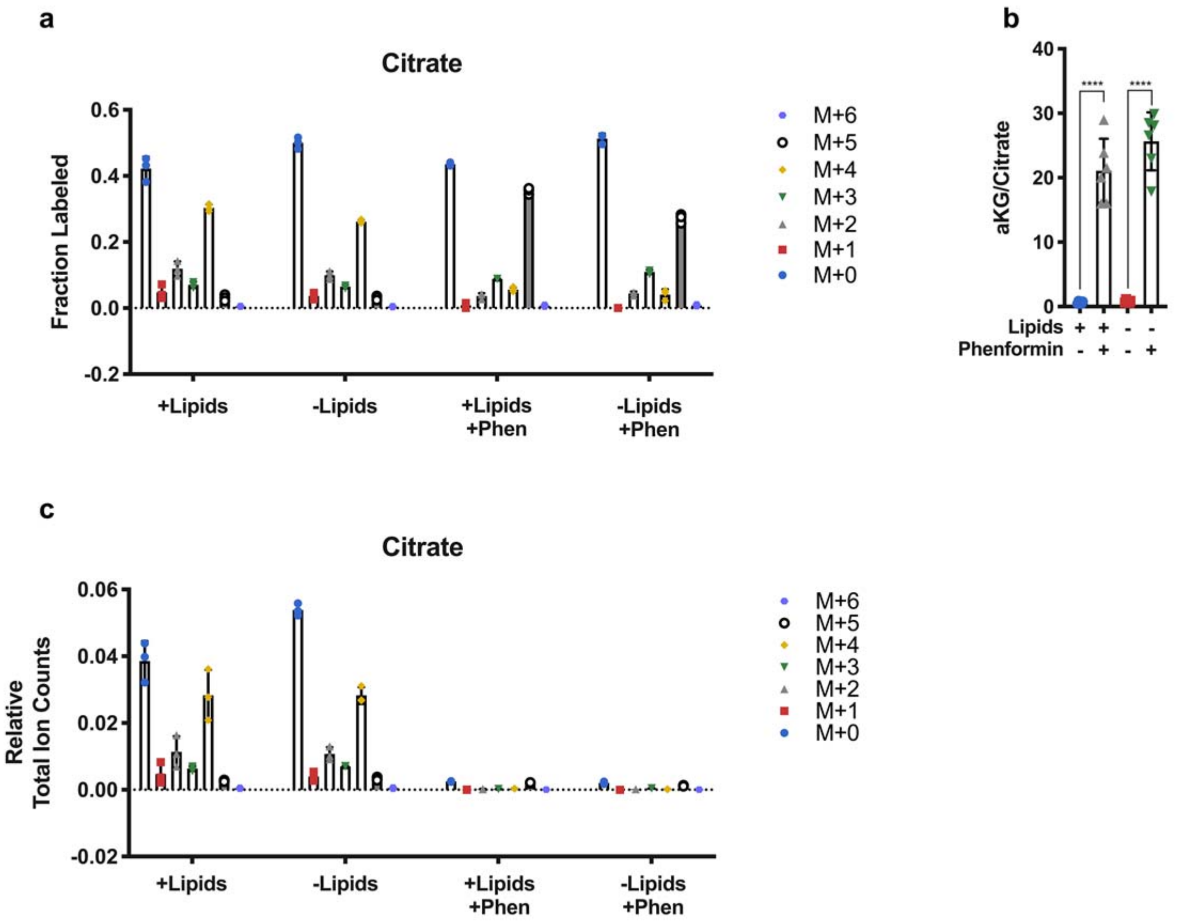


**Extended Data Fig. 4 | Inhibition of mitochondrial electron transport decreases intracellular citrate levels. a**, Relative fractional distribution of citrate isotopomers in HeLa cells cultured for 24 hours in media +lipids or –lipids with U-$^{13}$C-Glutamine, without and with and phenformin as indicated (100µM). **b**, Normalized intracellular ratio of αKG to citrate in HeLa cells cultured in +lipids or –lipids with or without phenformin (100µM). (n=6 per condition from a representative experiment). All data represent mean ± s.d. ***$P$ < 0.001, unpaired Student's $t$-test. All experiments were repeated 3 times or more. **c**, Isotopomer distribution of total levels of intracellular citrate in HeLa cells cultured for 24 hours in media +lipids or –lipids with U-$^{13}$C-Glutamine, without and with phenformin as indicated (100µM).

(n=3 per condition from a representative experiment). (n=3 per condition from a representative experiment).

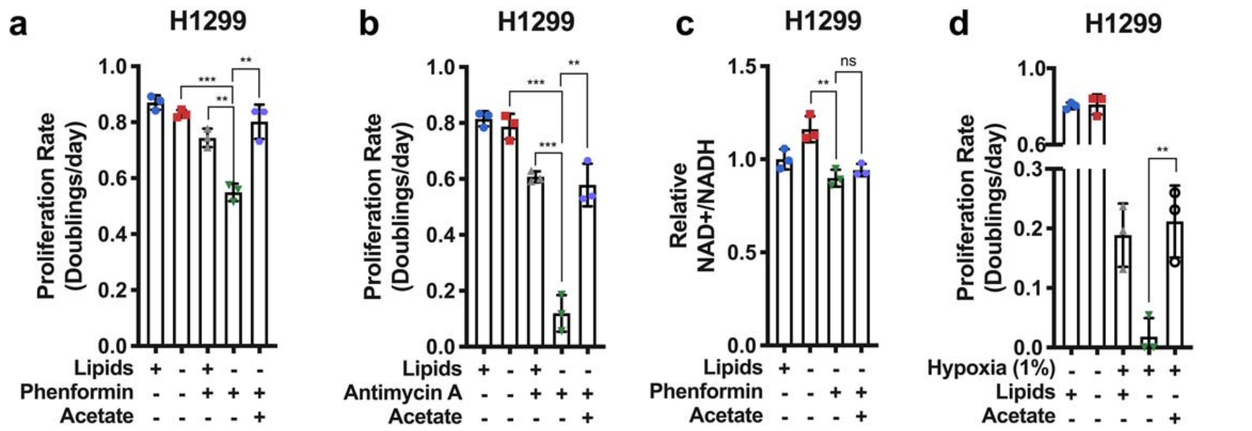


**Extended Data Fig. 5 | Bypassing oxidative steps in fatty acid synthesis rescues proliferation in electron acceptor deficient cells. a**, Proliferation rates of H1299 cells cultured in media +lipids or –lipids without or with phenformin (10µM) and/or acetate (200 µM) as indicated (n=3 per condition from a representative experiment). **b**, Proliferation rates of H1299 cells cultured in media +lipids or –lipids without or with antimycin A (15nM) and/or acetate (200 µM) as indicated (n=3 per condition from a representative experiment). **c**, Relative NAD+/NADH ratio in H1299 cells cultured in media +lipids or –lipids without or with phenformin (10µM) and/or acetate (200 µM) as indicated (n=3 per condition from a representative experiment). **d**, Proliferation rates of H1299 cells cultured in media +lipids or –lipids in normoxia (21% oxygen), hypoxia (1% oxygen), and/or acetate (200 µM) as indicated. Data from the first four conditions are the same as those presented in Extended Data Fig. 2a. (n=3 per condition from a representative experiment). All data represent mean ± s.d. $**P < 0.01$, $***P < 0.001$, ns $P > 0.05$, unpaired Student's *t*-test. All experiments were repeated 3 times or more.

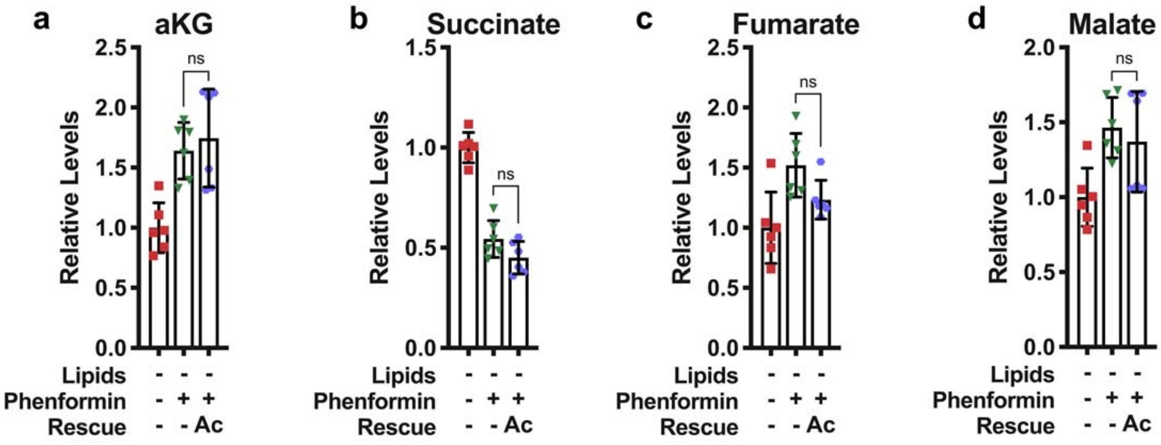


**Extended Data Fig. 6 | Effect of exogenous acetate on levels of TCA cycle intermediates. a**, Relative intracellular alpha-ketoglutarate levels in HeLa cells cultured for 24 hours in media –lipids without or with phenformin (100μM) and/or acetate (200 μM) as indicated (n=6 per condition from a representative experiment). **b**, Relative intracellular succinate levels in HeLa cells cultured for 24 hours in media –lipids without or with phenformin (100μM) and/or acetate (200 μM) as indicated (n=6 per condition from a representative experiment). **c**, Relative intracellular fumarate levels in HeLa cells for 24 hours in media –lipids without or with phenformin (100μM) and/or acetate (200 μM) (n=6 per condition from a representative experiment). **d**, Relative intracellular malate levels in HeLa cells cultured for 24 hours in media –lipids without or with phenformin (100μM) and/or acetate (200 μM) (n=6 per condition from a representative experiment). All data represent mean ± s.d., ns $P > 0.05$, unpaired Student's *t*-test. All experiments were repeated 3 times or more.

a

| TUMOR TYPES | FULL NAME | Correlation between hypoxia and lipid synthesis | p value (log 10) (red values are significant) | Correlation between hypoxia and lipid uptake | p value (log 10) (red values are significant) |
|---|---|---|---|---|---|
| ACC | Adenoid cystic carcinoma | -0.127 | -0.579 | 0.278 | -1.882 |
| BLCA | Bladder Urothelial Carcinoma | -0.474 | -23.474 | 0.176 | -3.456 |
| BRCA | Breast invasive carcinoma | -0.256 | -17.114 | 0.490 | -66.473 |
| CESC | Cervical squamous cell carcinoma and endocervical adenocarcinoma | -0.197 | -3.273 | -0.013 | -0.083 |
| CHOL | Cholangiocarcinoma | -0.430 | -2.051 | 0.341 | -1.377 |
| COAD | Colon adenocarcinoma | -0.366 | -9.733 | 0.375 | -10.231 |
| COADREAD | Colorectal adenocarcinoma | -0.382 | -13.853 | 0.405 | -15.664 |
| DLBC | Lymphoid Neoplasm Diffuse Large B-cell Lymphoma | -0.052 | -0.141 | 0.465 | -3.062 |
| ESCA | Esophageal carcinoma | -0.495 | -12.059 | -0.094 | -0.694 |
| GBM | Glioblastoma multiforme | -0.292 | -3.593 | 0.593 | -15.154 |
| GBMLGG | Glioma | -0.513 | -45.459 | 0.764 | -128.377 |
| HNSC | Head and Neck squamous cell carcinoma | -0.382 | -18.804 | 0.260 | -8.768 |
| KIPAN | Pan kidney (KICH+KIRC+KIRP) | -0.787 | -187.214 | 0.311 | -20.694 |
| LAML | Acute Myeloid Leukemia | -0.309 | -4.447 | 0.538 | -13.652 |
| LGG | Brain Lower Grade Glioma | -0.395 | -19.982 | 0.642 | -60.501 |
| LIHC | Liver hepatocellular carcinoma | -0.293 | -8.047 | 0.028 | -0.227 |
| LUAD | Lung adenocarcinoma | -0.291 | -10.748 | 0.119 | -2.158 |
| LUSC | Lung squamous cell carcinoma | -0.300 | -11.188 | 0.311 | -11.994 |
| MESO | Mesothelioma | -0.402 | -3.937 | 0.479 | -5.582 |
| OV | Ovarian serous cystadenocarcinoma | -0.261 | -5.379 | 0.422 | -13.778 |
| PAAD | Pancreatic adenocarcinoma | -0.466 | -10.240 | 0.294 | -4.166 |
| PCPG | Pheochromocytoma and Paraganglioma | -0.313 | -4.712 | 0.734 | -30.792 |
| PRAD | Prostate adenocarcinoma | -0.525 | -35.783 | 0.481 | -29.396 |
| READ | Rectum adenocarcinoma | -0.435 | -4.925 | 0.520 | -7.096 |
| SARC | Sarcoma | -0.253 | -4.434 | 0.437 | -12.788 |
| SKCM | Skin Cutaneous Melanoma | -0.385 | -4.225 | 0.436 | -5.379 |
| STAD | Stomach adenocarcinoma | -0.293 | -8.941 | 0.229 | -5.611 |
| STES | Stomach and Esophageal carcinoma | -0.351 | -18.064 | 0.074 | -1.142 |
| TGCT | Testicular Germ Cell Tumors | -0.290 | -3.499 | 0.304 | -3.801 |
| THCA | Thyroid carcinoma | -0.293 | -10.677 | 0.152 | -3.192 |
| THYM | Thymoma | -0.127 | -0.778 | 0.323 | -3.503 |
| UCEC | Uterine Corpus Endometrial Carcinoma | -0.209 | -2.273 | 0.403 | -7.551 |
| UCS | Uterine Carcinosarcoma | -0.362 | -2.243 | 0.417 | -2.898 |
| UVM | Uveal Melanoma | -0.121 | -0.548 | 0.318 | -2.397 |

b

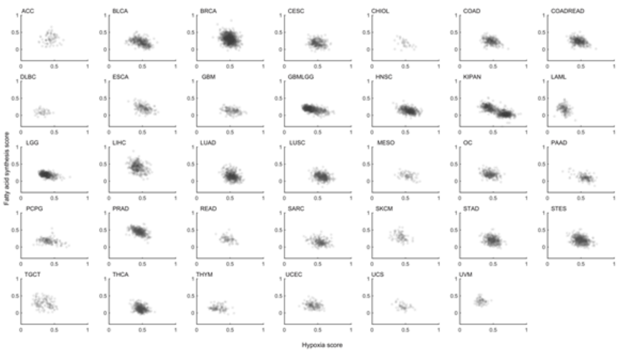


c

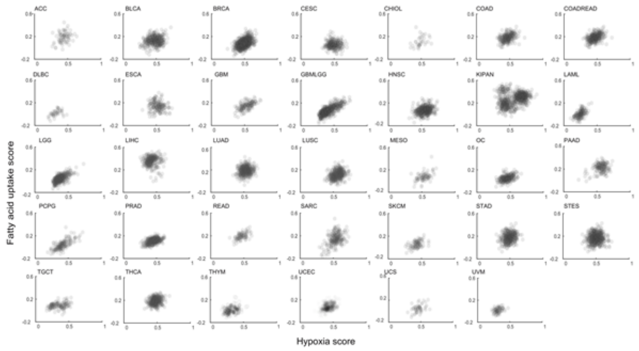


**Extended Data Fig. 7 | Gene expression correlations between lipid metabolism genes and hypoxia signature genes. a**, Pearson correlation coefficients, stratified by tissues of origin, between the tumor hypoxia score and expression of fatty acid uptake or fatty acid synthesis genes. All correlation p-values above the FDR=1% threshold are marked in red. **b**, Scatter plots across various tumor types show the correlations between the tumor hypoxia score and expression of genes participating in fatty acid synthesis. **c**, Scatter plots across various tumor types show the correlations between the tumor hypoxia score and expression of genes participating in fatty acid uptake.
